# Geometry-derived Hamiltonian organization distinguishes recurrent biological Fe–S architectures

**DOI:** 10.64898/2026.07.16.739031

**Authors:** Ji-Yong Sung, Jae-Ho Cheong

## Abstract

Iron–sulfur (Fe–S) clusters are among the most ancient biological cofactors, yet the physical properties associated with the recurrent biological utilization of specific Fe–S architectures remain unclear.

Here, we analyzed 2,404 experimentally resolved Fe–S clusters using an integrated framework combining structural geometry, geometry-derived effective-coupling network robustness, and coarse-grained open-system quantum transport simulations.

We show that 4Fe–4S clusters occupy a compact and partially distinct region of structural-descriptor space characterized by low geometric distortion, elevated geometry-derived network robustness, and broad taxonomic representation within the available structural dataset.

Geometry-derived coarse-grained effective Hamiltonian reconstruction and Lindblad simulations further showed architecture-dependent differences in Hamiltonian organization and simulated transport behavior under a common set of model assumptions.

Together, these findings establish a hierarchical comparative framework linking Fe–S geometry, effective Hamiltonian organization, simulated open-system transport, and broad taxonomic recurrence, and suggest that geometry-derived transport organization may represent one physical property contributing to the recurrent biological utilization of Fe–S architectures.

## Introduction

Iron–sulfur (Fe–S) clusters are among the most ancient molecular architectures in biology.^1^ Present in organisms across all domains of life, these inorganic cofactors mediate essential electron-transfer reactions that drive respiration, photosynthesis, nitrogen fixation, sulfur metabolism, DNA repair, and numerous other redox processes.^2^ Their remarkable evolutionary persistence has long suggested that Fe–S chemistry emerged early during the origin of life and became integrated into the core energetic machinery of living systems.^3^ Consistent with the iron–sulfur world hypothesis, modern biological Fe–S clusters may preserve structural solutions that originated from prebiotic iron–sulfur minerals.^4,5^ Yet despite decades of biochemical and structural investigation, the physical principles underlying the extraordinary evolutionary conservation of specific Fe–S architectures remain unclear.^6^

Among biological Fe–S cofactors, the 4Fe–4S cubane cluster occupies a uniquely prominent position.^7^ It is found in respiratory Complex I, photosynthetic reaction centers, ferredoxins, hydrogenases, nitrogenases, radical S-adenosylmethionine enzymes, and numerous additional redox proteins spanning phylogenetically distant organisms.^8^ Such widespread recurrence suggests that 4Fe–4S clusters possess properties extending beyond the functional requirements of individual proteins.^1^ Conventional explanations emphasize catalytic versatility, tunable redox potentials, and flexible ligand coordination. Although these features contribute to biological function, they do not explain why essentially the same structural architecture has been repeatedly retained throughout billions of years of evolution.

A fundamental constraint governing all biological electron transfer is distance-dependent quantum tunneling. ^9^ The Moser–Dutton framework demonstrated that electron-transfer rates can be predicted from a small number of physical parameters, including donor–acceptor distance, driving force, and reorganization energy, revealing that transport efficiency decreases exponentially with increasing distance.^9^ Consequently, biological electron-transfer systems are expected to evolve under strong geometric constraints. While this framework successfully explains electron-transfer kinetics between individual redox centers, it does not address a broader evolutionary question: why have certain Fe–S architectures repeatedly emerged as preferred structural solutions across unrelated biological systems? ^10^

One possible explanation is that evolution favors electron-transfer architectures that maximize robustness.^11,12^ Robust organization is a universal feature of biological systems, enabling functionality to persist despite structural perturbations, environmental fluctuations, and evolutionary change. If similar principles govern biological electron transfer, evolution should preferentially retain Fe–S geometries that maintain efficient transport across diverse structural contexts. Under this view, the structural diversity of Fe–S clusters should not be randomly distributed but organized into a landscape in which particular architectures occupy robust regions associated with favorable transport properties and long-term evolutionary persistence.

Despite extensive structural surveys and quantum chemical studies of individual Fe–S proteins, this hypothesis has never been examined systematically. Whether biological Fe–S clusters occupy distinct structural basins, whether these basins differ in transport robustness, and whether robustness is associated with evolutionary conservation remain unknown. Addressing these questions requires quantitative analysis across thousands of experimentally resolved structures rather than isolated investigations of individual proteins. ^13,14^

Here we analyzed 2,404 experimentally resolved Fe–S clusters using an integrated framework combining structural geometry, transport robustness, evolutionary conservation, and open-system quantum transport simulations. ^15^ We show that major Fe–S architectures occupy partially separable regions of structural-descriptor space and identify a compact region enriched in broadly distributed 4Fe–4S clusters. Geometry-derived coarse-grained Hamiltonian construction and Lindblad quantum transport simulations further reveal that conserved Fe–S geometries encode architecture-specific quantum transport through distinct Hamiltonian organizations.^16,17^ These results establish a hierarchical framework linking Fe–S geometry, Hamiltonian organization, quantum transport, and broad evolutionary recurrence, suggesting that Hamiltonian organization may contribute to the long-term persistence of biological Fe–S architectures.

### 4Fe–4S architectures preferentially occupy a distinct region of structural-descriptor space

To investigate whether biological Fe–S clusters exhibit global organizational principles beyond their local biochemical functions, we assembled a structurally curated dataset of 2,404 experimentally resolved Fe–S clusters spanning all major domains of life (**Fig. 1A**). The dataset comprised 2Fe–2S, 3Fe–4S, 4Fe–4S, and additional Fe–S architectures, providing a broad representation of naturally occurring redox cofactors.

**Figure 1.**
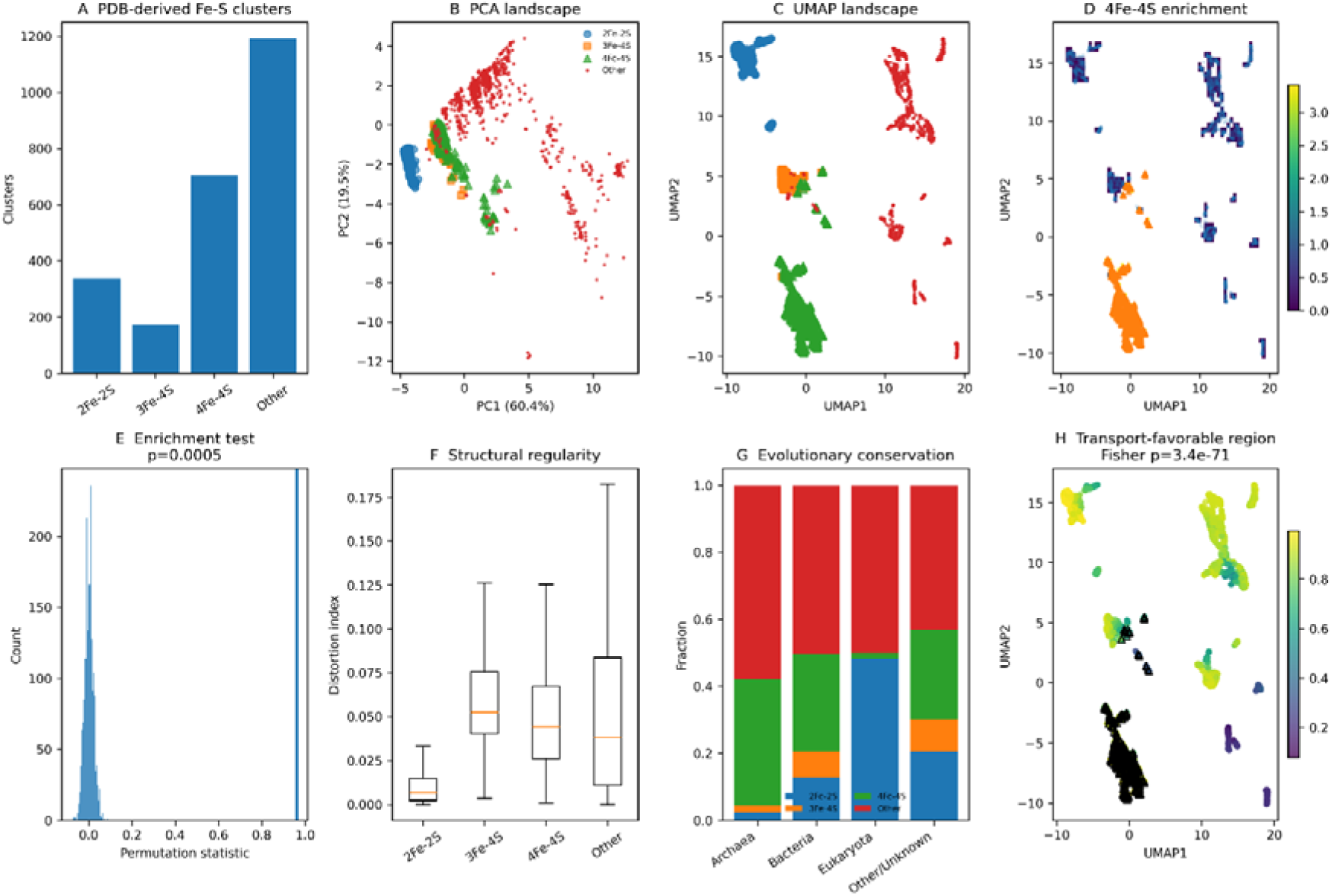
Structural landscape, evolutionary conservation, and transport-favorable regions of Fe–S clusters (A) Distribution of Fe–S cluster types identified from experimentally resolved protein structures in the Protein Data Bank (PDB). Clusters were classified into 2Fe–2S, 3Fe–4S, 4Fe–4S, and other Fe–S architectures. The majority of structures belong to the 4Fe–4S and other categories, providing a large structural dataset for comparative analysis. (B) Principal component analysis (PCA) of Fe–S cluster structural descriptors. Each point represents an individual Fe–S cluster projected onto the first two principal components. Distinct clustering patterns indicate that Fe–S architectures occupy separable regions of descriptor space. (C) Uniform Manifold Approximation and Projection (UMAP) embedding of the same descriptor set. The nonlinear embedding reveals discrete structural basins corresponding to different Fe–S cluster classes, suggesting the existence of preferred geometrical organizations. (D) Local enrichment of 4Fe–4S clusters across the UMAP landscape. Colors represent the local enrichment score calculated from neighborhood composition. Regions with elevated enrichment identify structural basins preferentially occupied by 4Fe–4S clusters. (E) Permutation-based enrichment analysis. The observed enrichment statistic (vertical line) is compared with the null distribution generated from randomized cluster assignments. The observed value lies outside the null distribution (*P* = 5 × 10), demonstrating significant enrichment of 4Fe–4S clusters within specific structural regions. (F) Structural regularity measured by the distortion index for each Fe–S cluster class. Lower values indicate more regular geometric organization. Boxplots show the median, interquartile range, and whiskers extending to 1.5× the interquartile range. Differences between cluster types indicate distinct structural constraints. (G) Evolutionary conservation across taxonomic domains. Stacked bars show the relative abundance of Fe–S cluster classes among major phylogenetic groups (Archaea, Bacteria, Eukaryota, and unclassified taxa). The widespread occurrence of 4Fe–4S clusters across diverse taxa suggests strong evolutionary retention of this architecture. (H) Transport-favorable regions within structural descriptor space. UMAP coordinates are colored according to a transport-favorability score derived from structural metrics associated with electron-transfer efficiency. Black triangles indicate 4Fe–4S clusters. Regions exhibiting high transport scores are significantly enriched in 4Fe–4S clusters (Fisher’s exact test, *P* = 3.4 × 10 ¹), suggesting that this architecture preferentially occupies structural configurations conducive to efficient electron transport.

We first quantified geometric variation among Fe–S clusters using a multidimensional descriptor set incorporating Fe–Fe distances, geometric regularity, compactness, and coordination statistics. Principal component analysis revealed substantial separation among major Fe–S architectures, indicating that cluster classes occupy distinct regions of structural space rather than forming a continuous geometric spectrum (**Fig. 1B**). In particular, 4Fe–4S clusters exhibited a compact distribution relative to other architectures, suggesting the existence of a highly conserved structural solution. To further characterize the global organization of Fe–S cluster space, we applied Uniform Manifold Approximation and Projection (UMAP) to the complete descriptor matrix. The resulting embedding revealed several discrete structural basins corresponding to different Fe–S architectures (**Fig. 1C**). Notably, 4Fe–4S clusters formed a dense and well-defined region that remained separated from both 2Fe–2S and 3Fe–4S clusters. This observation indicates that the cubane architecture occupies a distinct region of structural space rather than representing a gradual extension of other Fe–S topologies. We next asked whether this structural basin simply reflects the abundance of 4Fe–4S clusters or instead represents a statistically significant concentration within a preferred region of structural space. Local enrichment analysis demonstrated that the 4Fe–4S basin exhibited enrichment levels substantially exceeding global expectations (**Fig. 1D**). Permutation testing confirmed that this enrichment was highly unlikely to arise from random assignment of cluster identities (*P* = 5 × 10; **Fig. 1E**), indicating that the observed clustering reflects a genuine organizational feature of biological Fe–S architecture space. One possible explanation for this enrichment is that 4Fe–4S clusters occupy geometrically favorable configurations. Consistent with this hypothesis, 4Fe–4S clusters displayed significantly lower structural distortion than alternative Fe–S architectures (**Fig. 1F**). The reduced distortion of the cubane topology suggests enhanced geometric regularity and potentially more favorable electron-transfer pathways. Because evolutionary selection acts on function rather than geometry alone, we next examined the phylogenetic distribution of Fe–S architectures. Mapping cluster occurrence across Archaea, Bacteria, and Eukaryota revealed that 4Fe–4S clusters possess the broadest taxonomic representation among all major Fe–S classes (**Fig. 1G**).

This widespread representation indicates broad taxonomic recurrence across the sampled structural dataset and is consistent with the possibility that 4Fe–4S architectures confer a general functional advantage.

To determine whether the enriched structural basin is associated with electron-transfer performance, we calculated a transport-favorability score incorporating Fe–Fe spacing, geometric regularity, and cluster compactness. Projection of this score onto the structural landscape revealed a striking correspondence between transport-favorable regions and the 4Fe–4S-enriched basin (**Fig. 1H**). Regions exhibiting the highest transport-favorability scores were significantly enriched for 4Fe–4S architectures (Fisher’s exact test, *P* = 3.4 × 10 ¹), suggesting that the observed structural organization is linked to electron-transfer efficiency. Taken together, these results demonstrate that biological Fe–S clusters occupy a structured landscape rather than a continuous geometric continuum. Within this landscape, 4Fe–4S architectures form a statistically enriched basin characterized by low geometric distortion, favorable transport geometry, and broad evolutionary conservation. These findings suggest that the broad biological recurrence of 4Fe–4S clusters may be associated with the structural robustness of this electron-transfer architecture.

### Robust electron-transfer architectures emerge from higher-order Fe–S network topology

To determine whether the organization of Fe centres contributes to the robustness of Fe–S electron-transfer networks, we compared effective-coupling topologies across the major Fe–S architectures. Representative networks revealed a progressive increase in topological connectivity with cluster nuclearity: 2Fe–2S clusters formed a single effective Fe–Fe connection, 3Fe–4S clusters formed a triangular network, and 4Fe–4S clusters formed a densely interconnected topology containing multiple alternative coupling routes (**Fig. 2A**). This higher-order connectivity provides a structural basis for multiple potential effective-coupling routes and suggests that 4Fe–4S architectures may be less dependent on any single effective coupling. We next examined whether differences in network robustness could be attributed primarily to Fe–Fe geometry. Analysis of experimentally resolved PDB structures showed broadly overlapping Fe–Fe distance distributions across the major Fe–S architectures (**Fig. 2B**). Although the distributions differed statistically among groups (Kruskal–Wallis test, *P* < 10^−1^^6^), their absolute separation was modest, with median Fe–Fe distances remaining within a relatively narrow range. In particular, the Fe–Fe distances of 4Fe–4S clusters substantially overlapped those of 2Fe–2S, 3Fe–4S and other Fe–S architectures. Thus, variation in local Fe–Fe spacing alone is unlikely to account for the marked differences in network-level robustness. Because the number of possible Fe–Fe connections increases automatically with Fe-node number, direct comparison of raw network properties could confound higher-order geometric organization with the intrinsic effect of cluster unclarity. We therefore evaluated each experimentally observed Fe–S cluster relative to a cluster-specific ensemble of 250 randomized three-dimensional Fe configurations matched for both Fe-node number and mean pairwise Fe–Fe distance. These null configurations preserved cluster unclarity and overall geometric scale while disrupting the experimentally observed higher-order arrangement of Fe sites. Accordingly, positive null-adjusted scores indicate that an observed Fe–S geometry supports greater effective-coupling network robustness than expected for randomized geometries with the same number of Fe sites and the same mean Fe–Fe distance. The resulting geometry-specific robustness score was centred near zero for 2Fe–2S clusters, whereas 3Fe–4S and 4Fe–4S clusters exhibited positive node-adjusted scores (**Fig. 2C**).

**Figure 2.**
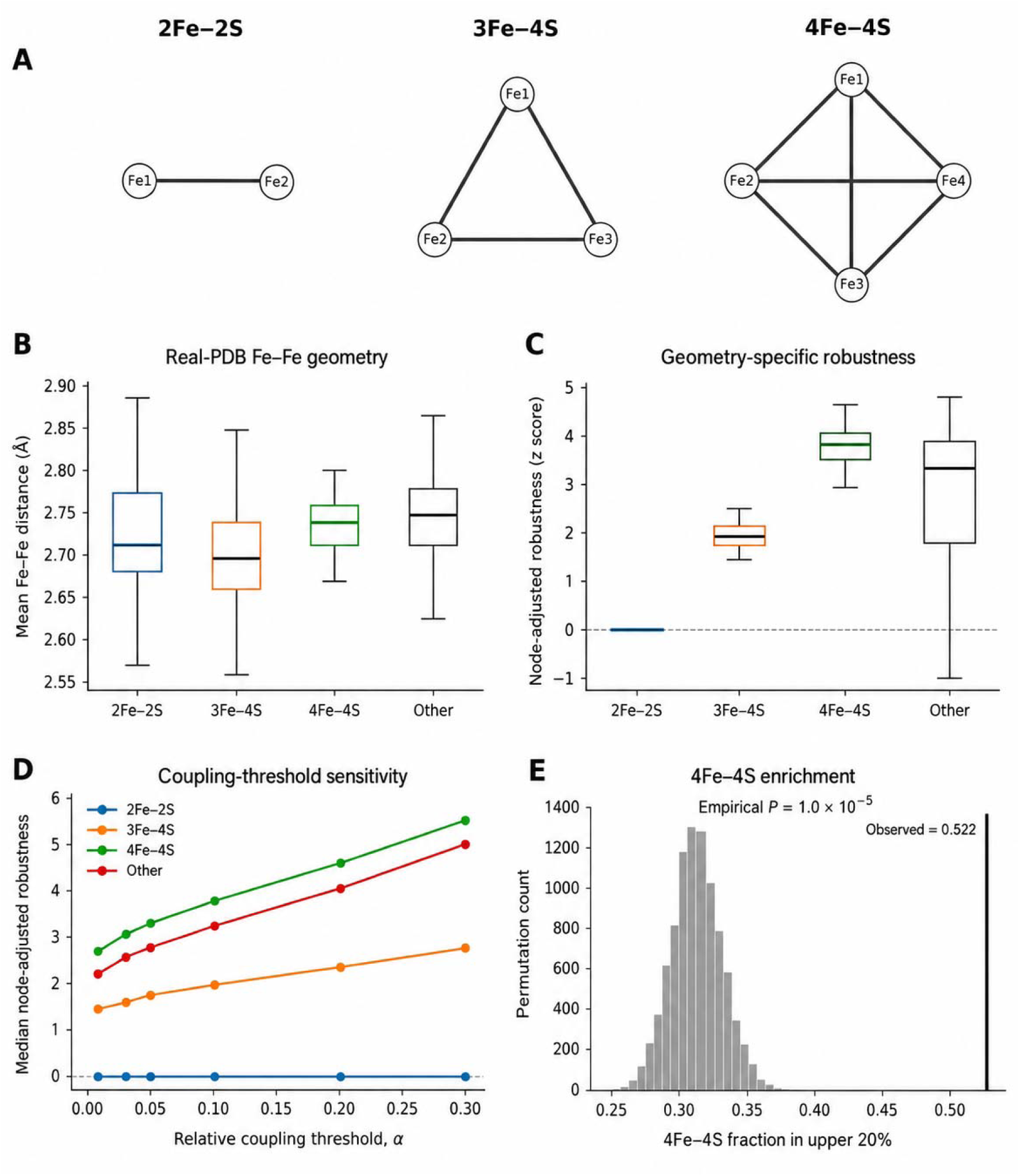
Geometry-specific robustness of biological Fe–S coupling networks (A) Schematic representations of effective electronic-coupling network topologies for 2Fe– 2S, 3Fe–4S and 4Fe–4S clusters. Nodes represent Fe sites and edges represent geometry-derived effective coupling between Fe sites. The diagrams illustrate network connectivity and are not crystallographic spatial projections. (B) Distribution of mean Fe–Fe distances calculated from experimentally determined PDB structures and grouped according to Fe–S cluster architecture. Box plots show the median, interquartile range (IQR) and whiskers extending to 1.5 × IQR; outliers are not shown. (C) Geometry-specific network robustness after correction using null networks matched for Fe-node number and geometric scale. Robustness values are expressed as null-adjusted z scores, with the dashed horizontal line indicating the null expectation (z = 0). Positive values indicate greater robustness than expected from the corresponding node-matched null model. (D) Sensitivity of median node-adjusted robustness to the relative effective-coupling threshold, α. The persistence of the relative ordering across the tested threshold range indicates that the observed architecture-dependent differences are not restricted to a single coupling cutoff. E, Enrichment of 4Fe–4S clusters among the upper 20% of node-adjusted robustness values. The histogram shows the null distribution generated by 10,000 random permutations of the architecture labels, and the vertical black line indicates the observed 4Fe–4S fraction (0.522). The empirical permutation test yielded P = 1.0 × 10.

Because 2Fe–2S clusters contain only a single Fe–Fe pair, their node- and scale-matched null configurations are geometrically degenerate; therefore, the near-zero adjusted score represents a reference baseline rather than evidence of intrinsically low robustness. Among the major canonical architectures, 4Fe–4S clusters showed the highest median geometry-specific robustness, with a median null-adjusted score of approximately 3.8, compared with approximately 2.0 for 3Fe–4S clusters. The broader distribution observed among other Fe–S architectures indicates substantial structural heterogeneity within this pooled category. Because each experimentally observed cluster was compared with randomized configurations matched for both Fe-node number and mean Fe–Fe distance, the elevated adjusted robustness of 4Fe–4S clusters cannot be explained solely by their larger number of Fe centers or by differences in overall geometric scale. Instead, the result reflects how the experimentally observed cubane-like arrangement organizes relative effective couplings, network connectivity, and alternative coupling routes compared with randomized geometries under the same matching constraints. We then tested whether this architectural advantage was dependent on the coupling threshold used to define effective network edges. The complete null-adjustment procedure was repeated at each threshold so that the observed geometries were evaluated relative to node- and scale-matched randomized configurations under the same network-construction criterion. Across the full range of relative coupling thresholds examined (α = 0.01–0.30), the ordering of the major Fe–S architectures remained stable (**Fig. 2D**). The median node-adjusted robustness of 4Fe–4S clusters remained consistently higher than that of 3Fe–4S and 2Fe–2S clusters and increased progressively as the coupling criterion became more stringent. The persistence of this relationship across threshold choices indicates that the observed 4Fe–4S advantage is not attributable to a single arbitrary network-construction parameter.

Finally, we tested whether the association between 4Fe–4S architecture and high node-adjusted robustness exceeded that expected from architecture-label assignment alone. Architecture labels were randomly permuted while the observed network properties were retained, generating a null distribution for the fraction of 4Fe–4S clusters within the upper 20% of the adjusted-robustness distribution. The observed fraction was 0.522, substantially exceeding the centre of the permutation distribution, which was approximately 0.32 (**Fig. 2E**). None of the 10,000 randomized assignments produced enrichment as large as that observed, yielding an empirical P value of 1.0 × 10^−4^. Thus, 4Fe–4S architectures are preferentially represented among highly robust networks beyond the expectation generated by randomized architecture labels.

Collectively, these analyses identify higher-order Fe-site organization as an important determinant of geometry-derived effective-coupling network robustness. The 4Fe–4S architecture retained the highest adjusted robustness after comparison with randomized geometries matched for Fe-node number and mean Fe–Fe distance, and this ordering remained stable across a broad range of relative coupling thresholds. These results indicate that the observed 4Fe–4S advantage is not explained solely by greater cluster unclarity or overall geometric scale. Rather, within the present network model, the cubane-like arrangement distributes effective coupling across multiple alternative routes and reduces dependence on individual connections. The present analysis does not directly demonstrate evolutionary selection or electron-transfer kinetics but identifies an architecture-dependent network property that may have contributed to the recurrent biological utilization of 4Fe–4S clusters.

### Robust Fe–S network architectures are distributed across taxonomic domains

To determine whether the robustness landscape identified from Fe–S network properties was restricted to a particular taxonomic lineage, we integrated PDB-derived structural data with taxonomic annotations. The resulting dataset spanned Archaea, Bacteria and Eukaryote, although representation was uneven across domains and a substantial fraction of structures lacked resolved domain-level assignments (**Fig. 3A**). Bacterial structures constituted the largest component of the dataset, whereas eukaryotic Fe–S clusters were comparatively under-represented. The fraction of clusters assigned as 4Fe–4S also varied among the sampled domains, with the highest proportion observed in Archaea. These distributions therefore describe the taxonomic coverage of available PDB structures rather than the biological prevalence of Fe–S architectures.

**Fig. 3.**
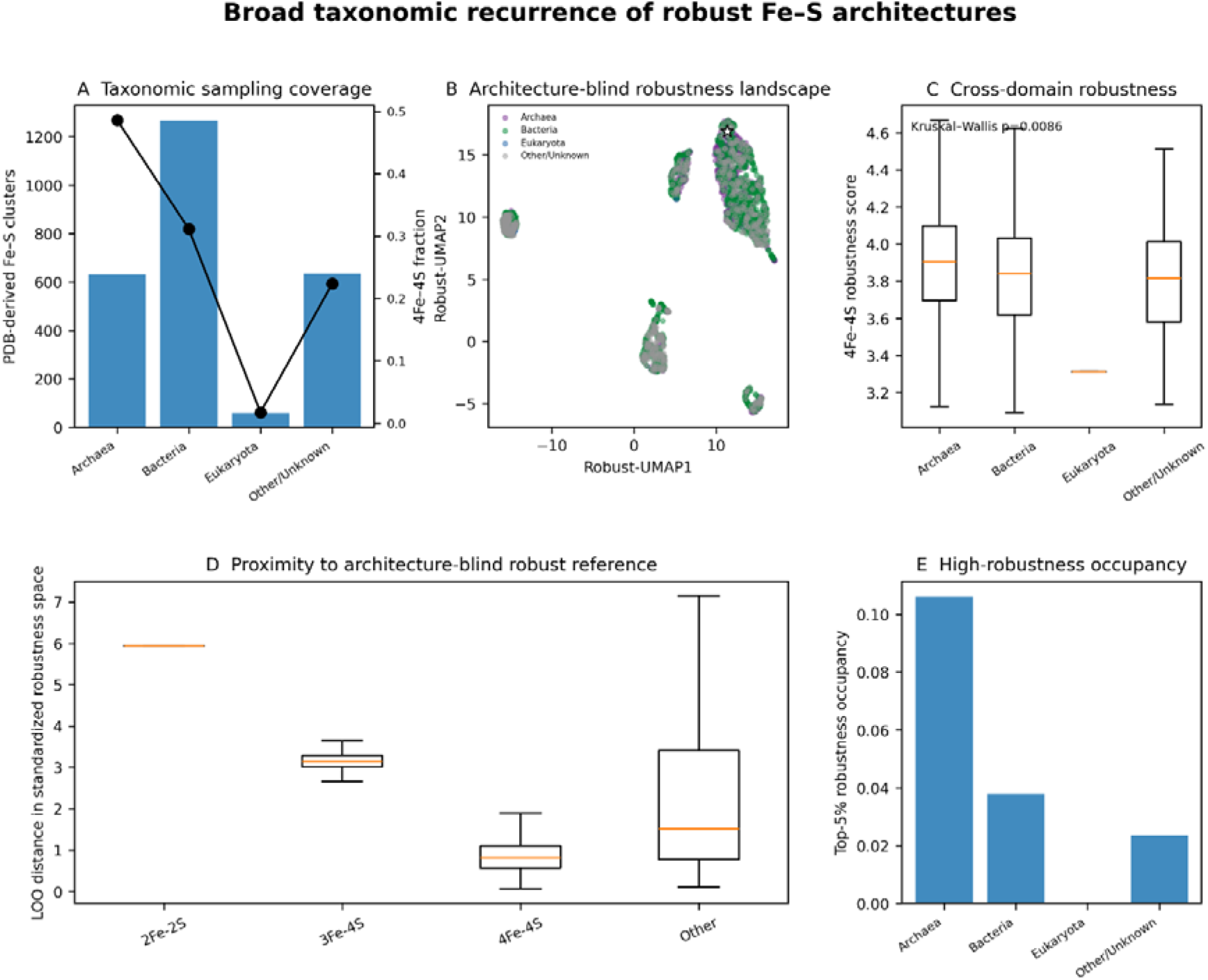
Architecture-blind analysis identifies robust Fe–S networks across taxonomic domains (A) Taxonomic composition of the PDB-derived Fe–S structural dataset. Bars indicate the number of Fe–S clusters assigned to each taxonomic domain (left axis), and the overlaid line indicates the fraction classified as 4Fe–4S within each domain (right axis). These values describe the taxonomic coverage of the available structural dataset and are not estimates of the biological prevalence of Fe–S architectures. (B) Architecture-blind robustness landscape generated by UMAP projection of standardized, node-adjusted network-robustness descriptors. Each point represents an individual PDB-derived Fe–S cluster and is coloured according to taxonomic domain. Architecture labels were not used to generate the robustness embedding or to define the high-robustness reference. The star indicates the centroid of the architecture-blind high-robustness reference set. UMAP coordinates are shown for visualization only; quantitative proximity was calculated in the original standardized robustness-feature space. (c) Distribution of node-adjusted robustness scores for 4Fe–4S clusters across sampled taxonomic domains. Box plots show the median (centre line), interquartile range (box) and values extending to 1.5 times the interquartile range (whiskers). Differences among domains were evaluated using a two-sided Kruskal–Wallis test (P = 0.0086). The Eukaryota category is represented by a single observation and is shown descriptively. (D) Leave-one-out distance of Fe–S clusters from the architecture-blind high-robustness reference in the original standardized robustness-feature space. For observations included in the reference set, the evaluated cluster was excluded before recalculation of the reference centroid. Lower distances indicate greater similarity to the multivariate robustness profile of the architecture-blind reference. Box plots show the median (centre line), interquartile range (box) and values extending to 1.5 times the interquartile range (whiskers). (E) Domain-specific occupancy of the upper 5% of the architecture-blind robustness distribution. Bars indicate the fraction of Fe–S clusters within each sampled taxonomic domain assigned to the high-robustness set. Occupancy values reflect the composition and unequal taxonomic representation of the PDB-derived structural dataset and should not be interpreted as estimates of evolutionary or biological prevalence.

We next constructed an architecture-blind robustness landscape using standardized, node-adjusted network descriptors. Fe–S architecture labels were excluded from both the dimensionality-reduction procedure and the definition of the high-robustness reference, thereby allowing robust network organization to be identified independently of prior structural classification. Projection of the resulting robustness-feature space revealed multiple regions occupied by clusters from different taxonomic domains (**Fig. 3B**). Archaea, Bacteria and Eukaryote showed substantial overlap within the landscape rather than forming completely domain-specific regions. The architecture-blind high-robustness reference localized to a region containing structures from multiple taxonomic backgrounds, indicating that the underlying robustness profile was not restricted to a single sampled domain. Because UMAP preserves local relationships but can distort global distances, the embedding was used only for visualization, whereas quantitative comparisons were performed in the original standardized robustness-feature space. To assess whether the robustness of 4Fe–4S networks was maintained across taxonomic groups, we compared node-adjusted robustness scores among sampled domains. 4Fe–4S clusters exhibited similarly elevated median robustness scores in Archaea, Bacteria and the Other/Unknown category, although the distributions differed significantly overall (two-sided Kruskal–Wallis test, *P* = 0.0086; **Fig. 3C**). The eukaryotic category contained only a single 4Fe–4S observation and was therefore considered descriptive rather than sufficient for domain-level inference. Thus, within the limits of the available structural sampling, elevated 4Fe–4S robustness was observed across multiple taxonomic backgrounds rather than being confined to one well-represented lineage. We then tested which Fe–S architectures most closely resembled the architecture-blind high-robustness state. To avoid circularity, proximity was quantified in the original standardized robustness-feature space using a leave-one-out procedure, in which each reference observation was excluded before recalculation of the reference centroid. 4Fe–4S clusters showed the shortest median distance to the architecture-blind robust reference, whereas 3Fe– 4S and 2Fe–2S clusters were progressively more distant (**Fig. 3D**). The broader distribution observed for other Fe–S architectures reflected the structural and topological heterogeneity of this pooled category. The close proximity of 4Fe–4S clusters emerged despite the exclusion of architecture labels from reference construction, indicating that the characteristic network properties of 4Fe–4S clusters independently converge on the multivariate robustness profile identified from the data.

Finally, we quantified the representation of each taxonomic domain within the upper 5% of the architecture-blind robustness distribution. High-robustness occupancy was greatest among archaeal structures, followed by bacterial and unclassified structures, whereas no eukaryotic clusters occurred within the upper 5% under the present sampling scheme (**Fig. 3E**). However, these occupancy values are influenced by the unequal representation and differing Fe–S architecture composition of the PDB-derived dataset and should not be interpreted as direct estimates of evolutionary frequency or biological prevalence. Instead, they demonstrate that highly robust Fe–S network states are represented across more than one sampled taxonomic domain. Together, these analyses show that the robust network organization associated with 4Fe–4S clusters is detectable without using architecture labels and is observed across multiple taxonomic backgrounds. The preferential proximity of 4Fe– 4S clusters to an independently defined high-robustness reference supports the interpretation that their densely interconnected topology occupies a distinct region of Fe–S network-property space.

These findings extend the null-adjusted analysis in Fig. 2 by showing that the 4Fe–4S robustness signature is not solely attributable to Fe-node number, mean geometric scale, explicit use of architecture labels, or a single dominant sampled taxonomic group. Rather than establishing independent evolutionary convergence directly, the results identify a broadly distributed network property that may have contributed to the recurrent biological utilization of 4Fe–4S clusters.

### Functional deployment of robust Fe–S network states

Having established that robust Fe–S architectures recur across broad taxonomic groups, we next asked whether these network properties are preferentially associated with specific biological functions. Fe–S-containing proteins were assigned to major functional classes using structure-linked annotations, and the distribution of Fe–S architectures and node-adjusted robustness was compared across respiratory/energy metabolism, biosynthesis/cofactor assembly and redox/electron-transfer functions (**Fig. 4**). The architectural composition of Fe–S clusters differed across functional contexts (**Fig. 4A**). Respiratory and energy-converting proteins showed the greatest representation of 4Fe–4S clusters, whereas biosynthetic/cofactor-assembly and redox/electron-transfer proteins contained more heterogeneous mixtures of Fe–S architectures. Consistent with this pattern, the fraction of clusters assigned as 4Fe–4S was highest in respiratory/energy-associated proteins, reaching approximately 0.37, compared with approximately 0.27 in biosynthesis/cofactor-assembly proteins and 0.28 in redox/electron-transfer proteins (**Fig. 4B**). These observations suggest that 4Fe–4S architectures are differentially deployed across functional environments rather than being uniformly distributed among Fe–S-dependent processes. We next examined whether functional deployment was associated with differences in network robustness. Respiratory/energy-associated Fe–S clusters displayed the highest median node-adjusted robustness, whereas biosynthesis/cofactor-assembly and redox/electron-transfer proteins exhibited lower median values and broader robustness distributions (**Fig. 4C**). Because node-adjusted robustness controls for the intrinsic effect of Fe-node number, this pattern cannot be attributed solely to the greater number of Fe centres in 4Fe–4S clusters. Instead, the results indicate that respiratory and energy-converting systems preferentially contain Fe–S configurations with comparatively robust network organization. To determine whether this association extended specifically to the most robust region of Fe–S architecture space, we quantified the occupancy of architecture-blind high-robustness states. High-robustness states were defined independently of cluster identity using the upper 5% of the node-adjusted robustness distribution. Respiratory/energy-associated clusters exhibited the highest occupancy of these states, with approximately 7.3% of clusters located within the high-robustness reference region (**Fig. 4D**). By comparison, high-robustness occupancy was approximately 0.9% for biosynthesis/cofactor-assembly proteins and 2.6% for redox/electron-transfer proteins. This preferential association remained evident after aggregation at the PDB-entry level, indicating that the pattern was not explained solely by the presence of multiple Fe–S clusters within individual structures. These results reveal a non-random relationship between biological function and Fe–S network robustness. Respiratory and energy-converting systems preferentially deploy highly robust Fe–S network states, extending the structural and taxonomic patterns identified in Figs. 2 and 3 into a functional context. Rather than demonstrating enrichment of a single Fe–S architecture alone, the results support a broader model in which biological energy-conversion systems preferentially occupy robust regions of Fe–S network space. Such organization may provide a structural basis for maintaining reliable electron-transfer connectivity under local geometric or network perturbations.

**Figure 4.**
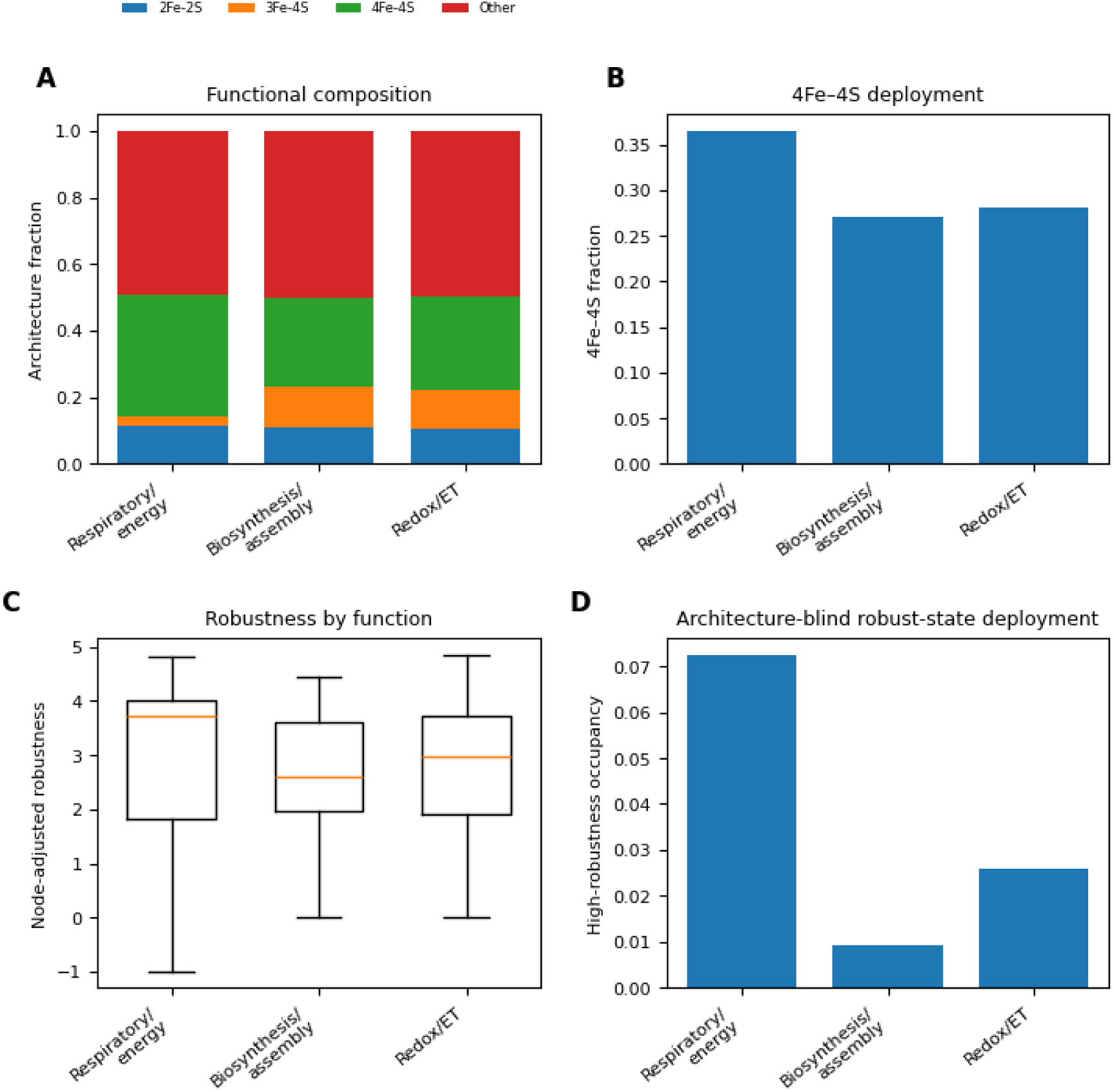
Functional deployment of robust Fe–S network states. (A) Functional composition of Fe–S cluster architectures across respiratory/energy metabolism, biosynthesis/cofactor assembly and redox/electron-transfer (Redox/ET) functional classes. Stacked bars indicate the relative fractions of 2Fe–2S, 3Fe–4S, 4Fe–4S and other Fe–S architectures within each functional class. (B) Fraction of Fe–S clusters assigned as 4Fe–4S within each functional class. Respiratory/energy-associated proteins showed the highest representation of 4Fe–4S clusters. (C) Distribution of node-adjusted network robustness across functional classes. Node-adjusted robustness quantifies geometry-associated network resilience after accounting for the effect of Fe-node number. Box plots show the median, interquartile range and whiskers extending to 1.5× the interquartile range; outliers are not shown. (D) Occupancy of architecture-blind high-robustness states across functional classes. High-robustness states were defined independently of Fe–S architecture using the upper 5% of the node-adjusted robustness distribution. Respiratory/energy-associated Fe–S clusters exhibited the highest occupancy of high-robustness network states. Together, these analyses indicate that robust Fe–S network configurations are preferentially deployed in respiratory and energy-converting systems rather than being uniformly distributed across functional contexts.

### Fe–S architectures generate distinct transport behavior in a coarse-grained open-system model

To investigate how Fe–S cluster geometry influences quantum electron transport, we reconstructed atomically resolved Hamiltonians from experimentally determined Fe–S structures and analyzed open-system transport across 1,266 Fe–S clusters. Each Fe atom was represented as a quantum transport site, while pairwise electronic couplings were parameterized using a distance-dependent tunneling model derived from Fe–Fe atomic coordinates. Lindblad master-equation simulations were then used to quantify transport efficiency under environmentally coupled quantum dynamics (**Fig. 5A**).

**Figure 5.**
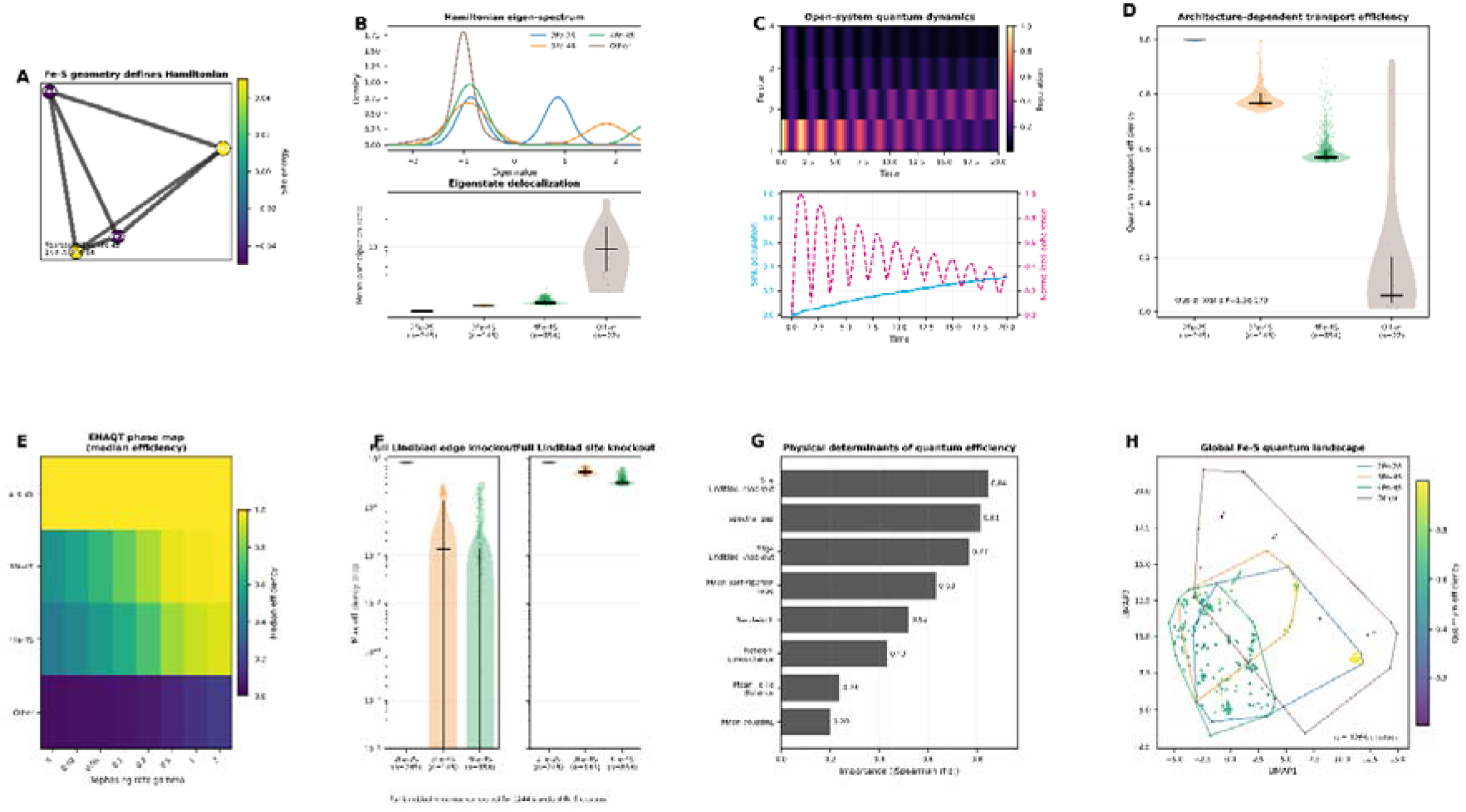
Fe–S cluster geometry encodes Hamiltonian properties that determine quantum transport efficiency and environment-assisted optimization. (A) Representative 4Fe–4S cluster illustrating the atom-derived quantum Hamiltonian. Fe atomic positions define the electronic coupling network, where edge thickness represents the coupling strength derived from Fe–Fe distances and node color denotes the site energy used in the Hamiltonian. (B) Architecture-dependent Hamiltonian properties. The upper panel shows the distribution of Hamiltonian eigenvalue spectra across Fe–S architectures, while the lower panel summarizes the corresponding eigenstate delocalization quantified by the mean participation ratio. Distinct Fe–S architectures generate characteristic Hamiltonian landscapes with different degrees of electronic delocalization. (C) Open-system quantum dynamics of a representative Fe–S cluster simulated using the Lindblad master equation. The upper panel shows the spatiotemporal population evolution across Fe sites. The lower panel presents sink population (cyan) and normalized quantum coherence (magenta), demonstrating coherent excitation transport followed by irreversible trapping at the terminal acceptor. (D) Architecture-dependent quantum transport efficiency for all analyzed Fe–S clusters. Violin plots show the distribution of transport efficiencies for each structural class. Horizontal black bars indicate the median, and vertical bars indicate the interquartile range. Sample numbers are indicated below each architecture. Statistical significance was evaluated using the Kruskal–Wallis test. (E) Environment-assisted quantum transport (ENAQT) phase map. Median transport efficiency is shown as a function of dephasing rate (_γ_) for each Fe–S architecture. The results reveal architecture-specific responses to environmental noise and identify optimal dephasing regimes that maximize transport efficiency. (F) Global sensitivity of quantum transport to structural perturbations determined by full Lindblad simulations. Left, maximum reduction in transport efficiency following removal of individual electronic couplings (edge knockout). Right, maximum reduction following removal of individual Fe sites (site knockout). These analyses identify critical transport bottlenecks across all Fe–S architectures. (G) Physical determinants of quantum transport efficiency ranked according to the absolute Spearman correlation with transport efficiency. Hamiltonian-derived descriptors, including spectral gap, eigenstate delocalization, electronic coupling, and Lindblad knockout sensitivity, explain a substantial fraction of the observed variability in transport performance. (H) Global quantum transport landscape of all analyzed Fe–S clusters projected using UMAP. Each point represents one Fe–S cluster and is colored according to its quantum transport efficiency. Colored outlines indicate the structural architecture of each cluster, demonstrating that evolutionarily conserved Fe–S geometries occupy distinct regions of Hamiltonian space associated with characteristic transport efficiencies. Together, these analyses demonstrate that Fe–S cluster geometry determines the underlying quantum Hamiltonian, which governs coherent transport dynamics and enables environment-assisted optimization of electron transfer. The resulting Hamiltonian properties provide a unifying physical explanation for architecture-dependent quantum transport across naturally occurring Fe–S clusters.

The resulting Hamiltonians revealed that different Fe–S architectures generate distinct electronic landscapes. Hamiltonian eigenvalue distributions differed systematically among 2Fe–2S, 3Fe–4S and 4Fe–4S clusters, accompanied by architecture-dependent changes in eigenstate delocalization quantified by the participation ratio (**Fig. 5B**). These results indicate that atomic geometry directly determines the spectral organization of the underlying Hamiltonian, thereby defining the degree of quantum-state delocalization available for electron transport. Open-system simulations further demonstrated that these Hamiltonian differences produce distinct transport dynamics. Representative 4Fe–4S clusters exhibited coherent population propagation followed by irreversible trapping at the terminal sink, while quantum coherence decayed continuously during transport (**Fig. 5C**). Rather than remaining fully coherent or fully incoherent, electron transfer occurred within an intermediate dynamical regime in which coherent evolution and environmental decoherence jointly contributed to transport. Across the complete Fe–S dataset, transport efficiency was strongly stratified according to cluster architecture (**Fig. 5D**). This separation remained highly significant across all reconstructed structures (Kruskal–Wallis test), demonstrating that quantum transport efficiency is an intrinsic property of Fe–S structural organization rather than a consequence of individual cluster variability. Thus, recurrent Fe–S architectures encode reproducible geometry-derived Hamiltonian properties associated with distinct transport performance. Because biological electron transfer occurs in noisy environments, we next examined the influence of environmental dephasing. Moderate dephasing increased transport efficiency across multiple Fe–S architectures, whereas excessive dephasing progressively suppressed transport (**Fig. 5E**). This non-monotonic dependence is consistent with environment-assisted quantum transport (ENAQT), indicating that environmental fluctuations can enhance rather than simply disrupt electron transfer. Importantly, the optimal dephasing regime differed among Fe–S architectures, suggesting that structural organization modulates the balance between coherent dynamics and environmental relaxation. To identify structural determinants of efficient transport, we systematically evaluated the sensitivity of each reconstructed Hamiltonian to perturbations. Full Lindblad edge- and site-knockout simulations demonstrated that only a limited subset of couplings and Fe sites exerted disproportionately large effects on transport efficiency (**Fig. 5F**), indicating that quantum transport is governed by localized bottlenecks embedded within otherwise distributed electronic networks. These bottlenecks were conserved across many independent Fe–S structures, supporting the existence of common transport motifs. Correlation analysis further showed that Hamiltonian-derived descriptors—including spectral gap, eigenstate delocalization, electronic coupling strength and knockout sensitivity—were among the strongest predictors of transport efficiency (**Fig. 5G**). Collectively, these results show that transport efficiency is strongly associated with Hamiltonian-derived descriptors that integrate the electronic consequences of Fe–S geometry. Finally, projection of all geometry-derived Hamiltonians, into a low-dimensional manifold revealed distinct architecture-specific regions in quantum transport space (**Fig. 5H**). Clusters sharing the same structural architecture occupied similar regions of the landscape and exhibited comparable transport efficiencies, suggesting that recurrent Fe–S architectures are associated with characteristic Hamiltonian organizations and distinct quantum transport properties. These analyses establish a hierarchical modeling framework in which atomic Fe–S geometry defines a coarse-grained Hamiltonian organization, which in turn generates architecture-dependent open-system dynamics and simulated transport efficiencies. This establishes a comparative modeling framework linking Fe–S geometry to effective Hamiltonian organization and architecture-dependent transport behavior under a common set of coarse-grained physical assumptions.

## Discussion

The remarkable conservation of 4Fe–4S clusters across the tree of life has long suggested the existence of a fundamental evolutionary advantage, yet the physical basis of this persistence has remained unresolved.^18,19^ This work provides three conceptual advances toward addressing this question. First, we demonstrate that biological Fe–S clusters occupy a structured robustness landscape in which 4Fe–4S architectures preferentially localize within a distinct transport-favorable region rather than being randomly distributed throughout structural space.^19–21^ Second, we show that this structural organization is associated with broad evolutionary conservation and preferential deployment in biological processes requiring reliable long-range electron transfer, suggesting that transport robustness represents an important systems-level constraint on Fe–S architecture. Third, by constructing geometry-derived coarse-grained Hamiltonians and performing open-system quantum transport simulations, we establish a mechanistic framework linking Fe–S geometry, Hamiltonian organization, quantum transport dynamics, and evolutionary conservation. Together, these findings support the view that the broad biological recurrence of 4Fe–4S clusters may be associated with a structurally robust and transport-favorable organization.

A central result of this study is that Fe–S cluster architectures do not occupy structural space uniformly. Instead, dimensionality reduction analyses revealed discrete structural basins associated with specific cluster topologies. Among these, the 4Fe–4S cubane architecture forms a particularly compact and enriched region of structural space. This observation is noteworthy because conventional descriptions of Fe–S clusters typically emphasize local chemical properties, such as redox potential, ligand coordination, or catalytic function. Our results instead indicate that higher-order organizational principles emerge at the scale of the entire Fe–S architecture landscape. The existence of a discrete 4Fe–4S basin suggests that biological evolution repeatedly converges toward a restricted subset of feasible structural solutions rather than exploring all geometrically accessible configurations.

The robustness analyses provide a geometry-derived network explanation for the recurrent association of 4Fe–4S clusters with robust effective-coupling states. Importantly, robustness was not inferred from raw network connectivity alone. Because higher-unclarity clusters possess more possible Fe–Fe connections by construction, each experimentally observed geometry was evaluated relative to randomized three-dimensional configurations matched for both Fe-node number and mean Fe–Fe distance. The persistence of elevated 4Fe–4S robustness after this adjustment indicates that the result is not explained solely by the number of Fe sites or overall geometric scale. Instead, it reflects the organization of relative effective couplings within the experimentally observed cubane-like geometry.

These findings indicate that, within the present geometry-derived network model, the cubane-like arrangement supports multiple alternative effective-coupling routes and reduces sensitivity to the weakening or removal of individual connections.

Such redundancy is expected to be particularly advantageous in biological energy-conversion systems, where interruption of electron flow can have severe physiological consequences. Thus, robustness emerges as a plausible selective pressure linking molecular geometry to evolutionary fitness. These observations extend classical theories of biological electron transfer. The Moser–Dutton framework demonstrated that electron-transfer rates can be predicted from a small number of physical parameters, particularly donor–acceptor distance and driving force. In this view, biological electron-transfer systems are constrained by the physics of quantum tunneling and therefore require precise spatial organization. Our results suggest that these geometric constraints operate within a broader robustness landscape. Whereas the Moser–Dutton framework explains why particular distances support efficient electron transfer, the robustness landscape identified here may explain why specific network architectures are repeatedly selected over evolutionary time. In this sense, transport robustness represents a systems-level extension of classical electron-transfer theory. The evolutionary implications of these findings are particularly intriguing. The iron–sulfur world hypothesis proposes that Fe–S chemistry played a central role in the earliest stages of metabolic evolution. If modern Fe–S clusters indeed derive from ancient geochemical precursors, one might expect historical contingency to contribute substantially to their persistence. However, our analyses suggest that contingency alone cannot account for the observed patterns. Fe–S clusters sampled from archaeal, bacterial, and eukaryotic structures occupy overlapping regions of robustness space despite their broad taxonomic separation. This broad taxonomic recurrence is consistent with the possibility that robust transport organization contributed to the long-term retention of these architectures, although the present analysis cannot distinguish recurrent selection from inheritance through shared evolutionary ancestry. We therefore propose that the intrinsic robustness of the 4Fe–4S architecture may have contributed to its broad biological recurrence alongside historical and lineage-dependent factors. The functional analyses further support this interpretation.

4Fe–4S clusters are not distributed uniformly across biological processes but show increased representation in respiratory and energy-associated proteins within the present dataset. Moreover, functions exhibiting the strongest enrichment of 4Fe–4S architectures also tend to occupy highly robust regions of the transport landscape. These findings suggest that biological systems selectively deploy robust Fe–S architectures when reliable electron transfer becomes a dominant functional constraint. In this framework, robustness is not merely a structural property but a reusable biological design principle. More broadly, our results contribute to an emerging view that robustness constitutes a fundamental organizing principle of biological systems. Similar concepts have been identified in metabolic networks, developmental programs, ecological communities, and protein interaction networks. The present study extends this perspective to biological cofactors, demonstrating that robustness can be detected directly within the structural organization of electron-transfer architectures. Importantly, the robustness landscape identified here emerges without explicit optimization of function, suggesting that robust configurations represent natural attractors within Fe–S architecture space.

Several limitations should be considered. First, our analysis relies on experimentally resolved structures and therefore reflects currently available structural data rather than the complete diversity of biological Fe–S clusters. Because PDB representation is shaped by taxonomic, functional, and experimental sampling biases, structural prevalence should not be interpreted as a direct measure of evolutionary frequency. We therefore interpret taxonomic breadth as a proxy for broad evolutionary distribution rather than as a direct estimate of evolutionary conservation. Although the observed recurrence of transport-robust 4Fe–4S architectures across taxonomic domains is consistent with evolutionary convergence, explicit phylogenetic analyses and ancestral-state reconstruction will be required to distinguish recurrent selection from retention through shared evolutionary ancestry.

Second, the quantum transport analysis employed a geometry-derived coarse-grained Hamiltonian constructed directly from experimentally resolved Fe atomic coordinates. In this framework, electronic couplings were parameterized using distance-dependent effective interactions rather than explicit electronic-structure calculations. Consequently, the reconstructed Hamiltonians are intended to capture the topology and relative organization of biological electron-transfer networks rather than quantitatively reproduce the full electronic structure of individual Fe–S clusters. Nevertheless, applying an identical physically motivated coarse-grained Hamiltonian across all architectures enables systematic comparison of transport organization while avoiding architecture-specific parameter fitting. Future studies incorporating quantum chemical calculations, protein electrostatics, bridge-mediated superexchange, and atomistic molecular dynamics will further refine the physical realism of the Hamiltonian model. Although these descriptors capture key aspects of network organization, future work incorporating atomistic electron-transfer calculations, open quantum-system models, or molecular dynamics simulations may provide a more detailed mechanistic understanding. Third, the present study focuses on static structural representations and does not explicitly account for conformational dynamics, environmental fluctuations, or protein-specific regulatory effects. These limitations nevertheless point toward important future directions. Combining the robustness framework introduced here with quantum transport models, including Marcus-type hopping kinetics, Moser–Dutton electron-transfer theory, and open quantum-system approaches, may reveal how structural robustness translates into dynamical transport efficiency.

The present framework should be viewed as complementary to, rather than a replacement for, classical electron-transfer theories such as the Marcus formalism. Marcus theory provides a quantitative description of electron-transfer rates by incorporating reorganization energy, thermodynamic driving force, and electronic coupling for individual donor–acceptor pairs. In contrast, our objective was to investigate how the global structural organization of Fe–S architectures influences the robustness and efficiency of transport at the network level. By reconstructing architecture-derived Hamiltonians and evaluating open-system transport across a large structural dataset, we focus on organizational principles that extend beyond pairwise electron-transfer kinetics. Future integration of architecture-level robustness with Marcus-type hopping models and experimentally derived redox parameters will provide an important step toward quantitatively linking evolutionary optimization with biological electron-transfer kinetics. Similarly, extending the present analysis to respiratory complexes, photosynthetic reaction centers, hydrogenases, and nitrogenases may uncover common design principles governing biological energy transduction at larger organizational scales.

In conclusion, our analyses identify a previously unrecognized robustness landscape governing the organization of biological Fe–S clusters. Within this landscape, 4Fe–4S architectures occupy a transport-favorable region characterized by enhanced robustness, broad taxonomic recurrence, and preferential functional deployment. These findings suggest that the broad biological recurrence of 4Fe–4S clusters may partly reflect the functional advantages associated with a robust electron-transfer architecture. More generally, they establish a conceptual framework linking cofactor geometry, transport robustness, and evolutionary selection, providing a systems-level perspective on the architecture of biological redox chemistry.

## Methods

### Collection and curation of biological Fe–S cluster structures

Experimentally resolved iron–sulfur (Fe–S) cluster-containing protein structures were collected from the Protein Data Bank (PDB).^22^ Candidate structures were identified through automated screening of HETATM records and ligand annotations corresponding to Fe–S cofactors.^23^ Structures containing incomplete Fe coordinates, unresolved occupancies, or ambiguous cluster assignments were excluded.^22^ Fe–S clusters were classified according to their stoichiometry into four architectural categories: 2Fe–2S, 3Fe–4S, 4Fe–4S, and other Fe– S topologies.^19^ To minimize sampling bias arising from highly represented proteins, redundant structures were removed using a combination of PDB identifiers, cluster composition, and structural similarity criteria. For each cluster, Cartesian coordinates of all iron and inorganic sulfide atoms were extracted directly from atomic coordinate files. Clusters containing fewer than two iron atoms were excluded because electron-transfer topology cannot be meaningfully defined for isolated iron centers. The final curated dataset consisted of 2,404 unique Fe–S clusters spanning a broad phylogenetic and functional range.^23^

### Structural descriptor generation

To quantitatively characterize the geometric organization of biological Fe–S clusters, a comprehensive set of structure-derived descriptors was calculated for every curated cluster. ^24^ These descriptors were selected to capture complementary aspects of Fe–S architecture, including overall cluster size, geometric regularity, spatial compactness, and local coordination patterns, all of which are expected to influence electron-transfer organization.^25^ The Cartesian coordinates of all Fe atoms were extracted directly from experimentally resolved PDB structures. Pairwise Fe–Fe distances were calculated as 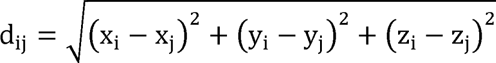, where x_i_, y_i_, and z_i_ denote the Cartesian coordinates of Fe atom i. From the pairwise distance matrix, a comprehensive set of structural descriptors was derived to quantify both global and local geometric properties of each Fe–S cluster. These descriptors comprised summary statistics of Fe–Fe distances (mean, minimum, maximum, and standard deviation), nearest-neighbor Fe–Fe distances, the radius of gyration,^26^ cluster compactness,^27^ coordination geometry statistics, and geometric distortion metrics. Collectively, these features provide a quantitative description of cluster size, symmetry, spatial organization, and packing density, thereby enabling systematic comparison of Fe–S architectures across the complete dataset.

The mean Fe–Fe distance was calculated as 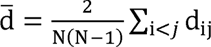 where *N* denotes the number of Fe atoms in the cluster. Cluster size was quantified using the radius of gyration, 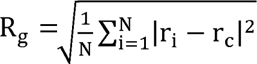 Where *r* represents the position vector of Fe atom i, and *r* denotes the centroid of all Fe atoms within the cluster. Smaller values of R_g_ indicate more spatially compact Fe–S architectures. To quantify structural regularity, a distortion index was defined as 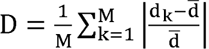 where d denotes an individual Fe–Fe distance, d is the mean Fe–Fe distance, and *M* is the total number of unique Fe–Fe pairs. Lower distortion values indicate highly symmetric architectures, whereas larger values reflect increased geometric irregularity. Cluster compactness was further evaluated as 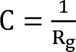 such that larger compactness values correspond to more densely packed Fe arrangements. Coordination geometry was characterized by the distribution of nearest-neighbor Fe–Fe distances together with local coordination statistics derived from the experimentally resolved atomic geometry. These descriptors quantify the spatial organization of neighboring Fe centers while remaining independent of cluster identity. Because the structural descriptors differ substantially in numerical scale and physical units, each descriptor was independently standardized prior to downstream analyses according to 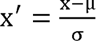 where µ and σ denote the mean and standard deviation calculated across the complete dataset. Standardization ensured that no individual descriptor dominated subsequent multivariate analyses because of its numerical magnitude rather than its structural information content. The resulting standardized descriptor matrix was used as the common input for principal component analysis (PCA), Uniform Manifold Approximation and Projection (UMAP), transport-favorability scoring, and robustness analyses.

### Principal component analysis

Principal component analysis (PCA) was performed as an unsupervised dimensionality-reduction method to identify the dominant sources of geometric variation among biological Fe–S clusters while preserving maximal variance within the original descriptor space.^28^ The covariance matrix of the standardized structural descriptor matrix was decomposed into orthogonal principal components by eigenvalue decomposition.^29^ Eigenvectors corresponding to the largest eigenvalues were retained, and each Fe–S cluster was projected onto the resulting principal component space using the associated loading vectors. The first two principal components captured the largest proportion of structural variance and were therefore used to visualize the global organization of Fe–S architectures. Because PCA was performed without incorporating cluster identity or functional annotations, the resulting representation provides an unbiased summary of intrinsic structural similarity among experimentally resolved Fe–S clusters.

### Uniform Manifold Approximation and Projection (UMAP)

To characterize nonlinear relationships among biological Fe–S architectures, Uniform Manifold Approximation and Projection (UMAP) was applied to the standardized structural descriptor matrix.^30^ UMAP was selected because it preserves local neighborhood relationships while maintaining the global organization of high-dimensional data, thereby providing an effective representation of continuous structural variation across diverse Fe–S clusters.

The embedding was generated using the parameters n_neighbors_ = 25, min_dist_ = 0.2, and the Euclidean distance metric. These parameters were chosen to balance preservation of local structural similarity with global manifold organization across the complete dataset. Each Fe– S cluster was projected into a two-dimensional embedding while retaining the neighborhood relationships present in the original descriptor space. Clusters located in close proximity within the UMAP embedding therefore exhibit similar geometric architectures, whereas distant clusters represent progressively distinct structural organizations. The resulting embedding provides a topology-preserving visualization of the structural landscape occupied by biological Fe–S clusters and forms the basis for all subsequent enrichment, robustness, and evolutionary analyses.

### Identification of 4Fe–4S-enriched structural basins

To determine whether 4Fe–4S architectures preferentially occupy specific regions of the structural manifold, local enrichment was quantified across the two-dimensional UMAP embedding using a two-dimensional grid-based density analysis.^31^ The embedding was partitioned into a uniform 70 x 70grid, and the number of all Fe–S clusters H_all_ and 4Fe– 4S clusters H_4Fe-4S_ within each grid cell were calculated independently. The local fraction of 4Fe–4S clusters was computed as 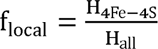 and normalized by the global fraction of 4Fe–4S clusters across the complete dataset, 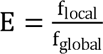 where f_global_ denotes the overall frequency of 4Fe–4S architectures.^32^ Grid cells containing fewer than three Fe–S clusters were excluded to reduce statistical noise arising from sparsely populated regions of the embedding. Grid cells exhibiting enrichment scores of E 1.5 were classified as enriched structural regions. Statistical significance of the observed enrichment was subsequently evaluated using permutation testing, in which 4Fe–4S labels were randomly shuffled while preserving the spatial distribution of the UMAP embedding.^33^

### Permutation-based significance testing

To determine whether the observed enrichment of 4Fe–4S architectures exceeded that expected by random spatial distribution, statistical significance was assessed using a permutation-based enrichment test.^31^ The two-dimensional UMAP embedding remained fixed throughout the analysis, while only the Fe–S cluster labels were randomly permuted, thereby preserving the overall abundance of each Fe–S architecture as well as the underlying geometric organization of the structural manifold. For each randomized dataset, local enrichment scores were recalculated using the identical grid-based enrichment procedure described above. A total of 3,000 independent random permutations were performed to generate the empirical null distribution of enrichment scores. The empirical significance of the observed enrichment was calculated as 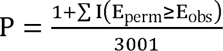 where E obs denotes the observed enrichment score, E_perm_represents enrichment scores obtained from randomized datasets, and I(.) is the indicator function. This permutation framework evaluates whether the observed localization of 4Fe–4S architectures within the structural manifold exceeds that expected from random assignment while preserving the global class composition of the dataset.

### Evolutionary conservation analysis

Taxonomic information associated with each experimentally resolved protein structure was obtained from NCBI taxonomy annotations together with RCSB structural metadata.^34^ Each Fe–S cluster was assigned to one of the three major domains of life (Archaea, Bacteria, or Eukaryote), enabling systematic comparison of phylogenetic distributions across all Fe–S architectures.^35^ Broad evolutionary distribution was approximated using taxonomic breadth, based on the premise that Fe–S architectures observed across diverse taxonomic lineages represent broadly distributed structural solutions.^36^ Taxonomic breadth was calculated as 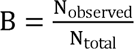 where N_observed_ denotes the number of distinct taxonomic lineages containing a given Fe–S architecture and N_total_ represents the total number of sampled taxonomic lineages within the curated dataset. Higher values of B therefore indicate broader representation across the sampled phylogenetic lineages, although this metric should not be interpreted as a direct measure of evolutionary conservation or evolutionary frequency.^37^ Taxonomic breadth was subsequently compared among Fe–S architectures to evaluate whether structurally enriched transport-favorable architectures also exhibited enhanced evolutionary persistence across the tree of life.

### Robustness metric justification and effective-coupling network construction

Electron transfer in biological Fe–S clusters requires the maintenance of effective electronic connectivity despite structural perturbations and local variations in Fe-site geometry. To characterize this architecture-level property, we employed graph-theoretical metrics that capture complementary aspects of effective-coupling network organization and perturbation tolerance. Each Fe atom was represented as a network node, and pairwise geometry-derived effective couplings between Fe sites were represented as weighted edges. Although this representation does not explicitly simulate quantum dynamics, it provides a coarse-grained description of effective-coupling organization and enables systematic comparison of network connectivity and robustness across structurally diverse Fe–S architectures. [38]

For both the experimentally observed and randomized Fe geometries, distance-dependent effective coupling weights were calculated using the same parameterization. To enable comparison across clusters, the effective couplings within each cluster were normalized relative to the strongest coupling in that cluster. An effective network edge was retained when its normalized coupling exceeded the predefined relative coupling threshold of α=0.10, which was used as the primary threshold throughout the analysis.

Five complementary network properties were calculated: normalized edge density, weighted global efficiency, bottleneck connectivity, edge-removal resilience, and normalized redundancy. Normalized edge density quantified the fraction of all possible Fe–Fe connections retained above the relative coupling threshold. Weighted global efficiency quantified network-wide effective connectivity using coupling-dependent shortest-path lengths. Bottleneck connectivity characterized the ability of the network to remain globally connected as the relative coupling threshold became progressively more stringent. Edge-removal resilience quantified the preservation of pairwise Fe-node connectivity during progressive removal of effective-coupling edges. Normalized redundancy quantified the availability of alternative cycle-forming connections after accounting for network size. Together, these descriptors capture complementary aspects of effective-coupling network organization, including connectivity, transport accessibility, bottleneck tolerance, perturbation resilience, and the availability of alternative coupling routes.

Because the number of possible pairwise Fe–Fe connections increases automatically with Fe-node number, direct comparison of raw network properties could intrinsically favor clusters of higher nuclearity. To account for this effect, each experimentally observed Fe–S cluster was evaluated relative to a cluster-specific geometric null ensemble. For each observed cluster, 250 randomized isotropic three-dimensional Fe configurations were generated using the same number of Fe nodes. Each randomized configuration was centered and uniformly rescaled so that its mean pairwise Fe–Fe distance matched that of the corresponding experimentally observed structure. The null configurations therefore preserved Fe-node number and overall geometric scale while disrupting the experimentally observed higher-order spatial arrangement of Fe sites.

For each network property m, the value obtained from the experimentally observed Fe geometry was standardized relative to the corresponding cluster-specific null distribution:

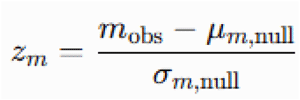

where m_obs_ denotes the network-property value calculated from the experimentally observed Fe geometry, and μ_m,null_ and σ_m,null_ denote the mean and standard deviation, respectively, obtained from the corresponding node- and scale-matched randomized geometries. When a null metric exhibited effectively zero variance, the corresponding adjusted value was assigned a neutral value of zero rather than interpreted as evidence of increased or decreased robustness.

The final geometry-specific robustness score was calculated as the arithmetic mean of the available component-specific null-adjusted z scores:

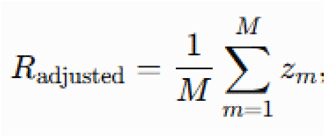

where M denotes the number of available robustness components. Positive values indicate that an experimentally observed Fe–S geometry exhibits greater effective-coupling network robustness than expected from randomized configurations with the same Fe-node number and mean pairwise Fe–Fe distance, whereas negative values indicate lower robustness than expected under the corresponding geometric null model. Because the null ensembles were matched for both Fe-node number and mean geometric scale, the adjusted score was used to distinguish architecture-dependent effective-coupling organization from the automatic increase in potential connectivity associated with higher cluster unclarity. For 2Fe–2S clusters, however, node- and scale-matched null configurations are geometrically degenerate because only one Fe–Fe pair exists. Accordingly, their adjusted score should be interpreted as a reference baseline rather than as evidence of intrinsically low robustness.

To evaluate whether the architecture-dependent robustness patterns depended on the effective-coupling cutoff used for network construction, the complete null-adjustment procedure was repeated across relative coupling thresholds ranging from α=0.01 to 0.30. Architecture-specific robustness patterns and rankings were compared across the complete threshold range. The persistence of the relative ordering among major Fe–S architectures was interpreted as evidence that the observed robustness differences were not restricted to a single arbitrary coupling threshold.

Importantly, these robustness metrics were designed to characterize geometry-derived effective-coupling organization and network perturbation tolerance rather than the kinetics of individual electron-transfer events. Unlike Marcus theory, which describes electron-transfer rates by incorporating electronic coupling, reorganization energy, and thermodynamic driving force, or Lindblad dynamics, which explicitly models open-system quantum transport under defined physical assumptions, the present graph-theoretical framework evaluates the architecture-level capacity of an effective-coupling network to maintain connectivity and alternative coupling routes under perturbation. The robustness framework therefore complements, rather than replaces, the explicit open-system quantum transport simulations described below.

### Node- and geometric-scale-adjusted robustness

Because the number of possible pairwise connections increases automatically with the number of Fe sites, direct comparison of raw network robustness across Fe–S architectures may favor clusters of higher unclarity. We therefore evaluated each experimentally observed Fe–S geometry relative to a cluster-specific geometric null ensemble matched for both Fe-node number and global spatial scale. For each observed cluster, 250 isotropic random three-dimensional point configurations were generated using the same number of Fe nodes. Each random configuration was centered and uniformly rescaled so that its mean pairwise Fe–Fe distance was identical to that of the corresponding experimental structure. This procedure preserved Fe-node number and mean geometric scale while disrupting the observed higher-order arrangement of Fe sites.

### Functional enrichment analysis

Proteins were assigned to functional categories using UniProt annotations and PDB metadata. ^38^ Categories included: respiratory and energy metabolism, oxidoreductases, ferredoxin/electron carriers, nitrogen and sulfur metabolism, photosynthesis, other enzymes. Enrichment of specific Fe–S architectures within functional groups was evaluated using Fisher’s exact test. Odds ratios and 95% confidence intervals were calculated from contingency tables. Multiple-testing correction was performed using the Benjamini– Hochberg procedure. Adjusted q values below 0.05 were considered statistically significant.^39^

### Statistical analysis

All statistical analyses were performed in Python. Continuous variables are reported as medians and interquartile ranges unless otherwise stated. Comparisons among multiple groups were performed using Kruskal–Wallis tests. Associations between continuous variables were quantified using Spearman rank correlation. All reported P values are two-sided unless otherwise indicated. Statistical significance was defined as *P* < 0.05.

### Geometry-derived Hamiltonian construction and open-system transport analysis

The structural analyses described above establish the geometric and topological organization of biological Fe–S clusters but do not directly reveal the physical mechanisms governing electron transport. To determine whether conserved Fe–S architectures encode distinct quantum transport properties, we constructed geometry-derived coarse-grained Hamiltonians directly from experimentally resolved Fe–S geometries and performed open-system quantum transport simulations for all Fe–S clusters included in the transport dataset.^15,39^ This framework provides a physically interpretable representation of architecture-dependent electronic organization while enabling systematic comparison across structurally diverse Fe– S clusters using a unified Hamiltonian formalism. By combining Hamiltonian reconstruction with Lindblad open-system dynamics, we evaluated how experimentally observed structural variation influences coherent transport, environmental decoherence, transport robustness, and overall quantum transport efficiency.^40,41^

### Atom-derived Hamiltonian construction

For each Fe–S cluster, the Cartesian coordinates of all Fe atoms were extracted from the curated structural dataset described above. Each Fe atom was represented as an effective quantum transport site because Fe centers constitute the primary redox-active components responsible for long-range electron transfer in biological Fe–S clusters. Sulfur atoms and the surrounding protein environment were not treated as explicit transport sites but were implicitly represented through the experimentally resolved Fe–Fe geometry encoded in the coarse-grained Hamiltonian. This representation preserves the overall topology of biological electron-transfer pathways while enabling systematic comparison across structurally diverse Fe–S architectures. We intentionally adopted a coarse-grained Hamiltonian to enable systematic comparison of thousands of experimentally resolved Fe–S clusters under identical physical assumptions rather than to reproduce the full ab initio electronic structure of individual cofactors. This strategy allows architecture-dependent differences in transport organization to emerge directly from experimentally observed structure al variation while avoiding cluster-specific parameter fitting. The electronic structure of each Fe–S cluster was represented by an effective single-particle tight-binding Hamiltonian, 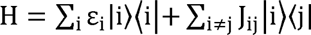 where ε_i_denotes the effective site energy associated with Fe site i, and J_ij_ represents the electronic coupling between Fe sites i and j. The first term describes the local electronic energy of each transport site, whereas the second term accounts for coherent electronic coupling between neighboring Fe centers.

In the present coarse-grained model, identical on-site energies were assigned to all Fe transport sites (ε_i_ = 0 for all i). This assumption was adopted to isolate the influence of geometry-dependent electronic couplings on Hamiltonian organization. Consequently, architecture-dependent differences in the Hamiltonian arise exclusively from the distance-dependent Fe–Fe electronic coupling matrix.

Electronic couplings were parameterized using an exponentially decaying distance-dependent interaction, 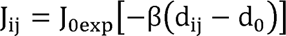 where d_ij_ is the experimentally observed Fe–Fe distance, (J_0) is the coupling strength at the reference distance d_0_, and β controls the decay of electronic coupling with increasing donor–acceptor separation. The exponential distance dependence reflects the well-established decay of biological electron-transfer coupling predicted by quantum tunneling theory and the Moser–Dutton relationship. The reference distance d_0_ and decay constant β were fixed throughout the analysis to ensure that all Fe–S architectures were evaluated using identical physical assumptions.

No architecture-specific fitting parameters were introduced. Consequently, differences in Hamiltonian spectra arise exclusively from experimentally observed structural variation rather than empirical model tuning, allowing unbiased comparison of electronic organization across all Fe–S architectures. The reconstructed Hamiltonians are intended to capture architecture-dependent electronic organization rather than reproduce the complete electronic structure of individual Fe–S clusters. This atom-derived coarse-grained representation provides a physically interpretable description of electron-transfer topology while remaining computationally tractable for large-scale comparative analyses. The resulting Hamiltonians served as the basis for all subsequent spectral characterization and open-system Lindblad transport simulations.

### Hamiltonian spectral characterization

Each reconstructed Hamiltonian was diagonalized using standard Hermitian eigensolvers implemented in SciPy to obtain the complete set of real-valued eigenenergies and orthonormal eigenvectors. These eigenstates provide a compact representation of the electronic organization encoded by each experimentally resolved Fe–S geometry. For every generated Hamiltonian, we calculated a set of complementary spectral descriptors, including the eigenvalue spectrum, spectral bandwidth, minimum eigenvalue gap, and mean participation ratio. The spectral bandwidth was defined as W = E_max_-E_min_ where E_max_ and E_min_ denote the largest and smallest Hamiltonian eigenvalues, respectively. The minimum eigenvalue gap was calculated as ΔE_min_ - min_i_ (E_i+1_-E_i_) providing a measure of the energetic separation between neighboring eigenstates. Eigenstate delocalization was quantified using the participation ratio, 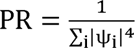 where ψ_i_ denotes the normalized amplitude of the eigenstate on Fe site i. Larger participation ratios indicate that quantum states are distributed across multiple Fe centers, whereas smaller values correspond to stronger spatial localization. These Hamiltonian-derived descriptors characterize complementary physical properties of the reconstructed electronic landscape, including spectral dispersion, energetic separation, and quantum-state delocalization. Together, they provide a physically interpretable representation of architecture-dependent electronic organization and form the basis for the comparative analyses presented in Fig. 5B as well as the subsequent quantum transport simulations.

### Lindblad quantum transport simulations

Quantum electron transport was simulated within the Markovian open quantum system formalism using the Lindblad master equation, which describes coherent Hamiltonian evolution together with irreversible environmental decoherence and excitation trapping. This framework was adopted because biological electron transfer occurs in fluctuating protein environments where coherent dynamics are continuously influenced by interactions with the surrounding molecular environment. The density matrix evolved according to 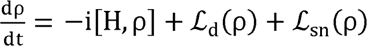 where the first term describes coherent evolution generated by the reconstructed Hamiltonian, whereas the Lindblad super operators account for pure dephasing and irreversible excitation trapping. The initial excitation was localized on the first Fe center, 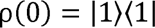 representing electron injection into the Fe–S cluster. Pure dephasing was independently applied to every Fe center using identical Lindblad operators, 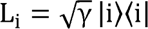 where γ denotes the environmental dephasing rate. Identical dephasing rates were used for all transport sites to isolate architecture-dependent transport properties from site-specific parameter variation.

Irreversible population trapping at the terminal Fe center was represented by a sink operator, 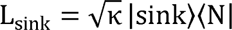 where κ denotes the trapping rate and *N* represents the terminal Fe transport site. This sink mimics irreversible electron transfer to the downstream electron acceptor. Time evolution was obtained by numerical exponentiation of the complete Liouvillian super operator until the final simulation time T. Quantum transport efficiency was defined as 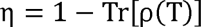 which corresponds to the fraction of excitation irreversibly transferred to the sink during the simulation. For representative Fe–S clusters, complete population dynamics and quantum coherence evolution were retained to generate the transport maps shown in Fig. 5C. The resulting transport efficiencies were subsequently compared across all reconstructed Hamiltonians to evaluate architecture-dependent quantum transport performance.

### Environment-assisted quantum transport (ENAQT)

To evaluate the influence of environmental noise on quantum transport, we performed environment-assisted quantum transport (ENAQT) simulations by systematically varying the environmental dephasing rate while keeping the reconstructed Hamiltonian unchanged. ENAQT predicts that intermediate levels of environmental dephasing can enhance transport efficiency by suppressing destructive quantum interference while preserving coherent transport pathways. For every reconstructed Fe–S Hamiltonian, the dephasing rate was varied over the range γ= 0, 0.01, 0.02, 0.05, 0.10, 0.20, 0.50, 1.0, 2.0 thereby spanning the transition from the fully coherent regime (γ = 0) to strongly environment-dominated transport. Throughout this analysis, all Hamiltonian parameters and sink dynamics were held constant so that changes in transport efficiency originated exclusively from environmental dephasing. For each Fe–S architecture, the transport efficiency was evaluated at every dephasing rate, and the median transport efficiency was calculated across representative structures of the same architecture. Median values were used because transport efficiencies exhibited non-Gaussian distributions with occasional extreme values. The resulting ENAQT profiles were used to construct the phase map shown in Fig. 5E and to identify architecture-dependent optimal dephasing regimes.

### Full Lindblad edge- and site-knockout analysis

To identify the structural elements that make dominant contributions to quantum transport, systematic edge- and site-knockout analyses were performed for every reconstructed Hamiltonian. These perturbation analyses quantify the sensitivity of quantum transport to the removal of individual electronic couplings and Fe transport sites. For edge-knockout simulations, each electronic coupling was individually removed by setting J_ij_ = 0 while preserving all remaining Hamiltonian elements. For site-knockout simulations, all electronic couplings associated with a selected Fe atom were simultaneously removed before repeating the complete Lindblad transport calculation, thereby preserving the remaining network topology while eliminating transport through the perturbed site. Transport sensitivity was quantified as Δη = η_original_ - η_knockout_ where larger values of Δη indicate greater dependence of quantum transport on the removed structural element. For every Fe–S cluster, the maximum transport reduction obtained following edge removal and site removal was retained as the global knockout sensitivity, thereby identifying the dominant transport bottleneck within each reconstructed Hamiltonian.

Unlike the graph-theoretical robustness analyses presented in Fig. 2, which evaluate network resilience using purely topological descriptors, the present knockout analyses explicitly recomputed the complete Lindblad quantum dynamics after every perturbation. Consequently, the reported sensitivities reflect the combined effects of Hamiltonian evolution, environmental decoherence, and irreversible excitation trapping on quantum transport efficiency. These architecture-specific knockout profiles constitute the quantum transport sensitivities presented in Fig. 5F.

### Identification of Hamiltonian determinants of transport efficiency

To identify the Hamiltonian properties most strongly associated with quantum transport performance, Hamiltonian-derived descriptors were systematically correlated with Lindblad transport efficiencies across all simulated Fe–S clusters. The evaluated descriptors were selected to capture complementary physical characteristics of the reconstructed Hamiltonians, including spectral organization, energetic dispersion, electronic-state delocalization, network connectivity, structural geometry, and transport robustness. The analyzed descriptors included the spectral gap, Hamiltonian bandwidth, mean participation ratio, mean electronic coupling, network conductance, mean Fe–Fe distance, maximum edge-knockout sensitivity, and maximum site-knockout sensitivity. Associations between each descriptor and transport efficiency were quantified using Spearman rank correlation coefficients because several Hamiltonian descriptors exhibited non-Gaussian distributions and potentially nonlinear monotonic relationships with transport performance. Absolute correlation coefficients were subsequently used to rank descriptor importance irrespective of whether individual descriptors exerted positive or negative effects on transport efficiency. The resulting ranking identifies the Hamiltonian features most strongly associated with architecture-dependent quantum transport performance and forms the basis of the determinant analysis presented in Fig. 5G.

### Hamiltonian landscape analysis

To investigate whether distinct Fe–S architectures also occupy organized regions in quantum transport space, Hamiltonian-derived descriptors and Lindblad transport properties were integrated into a unified descriptor matrix and projected into two dimensions using the same UMAP framework described for the structural analyses. The integrated descriptor matrix comprised Hamiltonian spectral descriptors, transport-efficiency metrics, ENAQT characteristics, and quantum robustness measurements, thereby providing a comprehensive representation of the electronic organization of each reconstructed Fe–S cluster. The same UMAP parameters used for the structural embedding were retained to facilitate direct comparison between geometric organization and Hamiltonian organization across the complete dataset. Whereas the structural embedding presented in Fig. 1 reflects similarities in experimentally resolved Fe–S geometry, the Hamiltonian landscape represents similarities in reconstructed electronic organization and quantum transport behavior. Each point in the resulting embedding corresponds to one reconstructed Hamiltonian, with neighboring points representing Fe–S clusters exhibiting similar Hamiltonian spectra, transport dynamics, and robustness characteristics. This Hamiltonian landscape therefore provides an integrated visualization of architecture-dependent quantum transport organization and forms the basis of the comparative analysis presented in Fig. 5H.

### Software implementation

All analyses were performed using custom Python scripts together with NumPy, pandas, SciPy, scikit-learn, UMAP-learn, and Matplotlib. All figures and statistical analyses were generated directly from the curated Fe–S cluster dataset without manual modification of individual observations.

## Declarations

### Ethics approval and consent to participate

N/A

### Consent for publication

N/A

### Availability of data and materials

The analysis codes and datasets used and/or generated during the current study are available from Dr. Ji-Yong Sung upon reasonable request. Interested researchers may contact Dr. Sung via email at to obtain access to the relevant materials.

### AI Use Declaration

AI tools were used only for English grammar correction and language polishing.

### Authors’ contributions

Conceptualization & Investigation: JYS, JHC; Methodology: JYS; Data analysis: JYS; Writing-original draft: JYS; Writing-review & editing: JYS, JHC; Supervision: JYS, JHC; Project administration: JYS, JHC; Funding acquisition: JHC; Interpretation of the results; JYS, JHC. All authors have read and agreed to the published version of the manuscript.

## Acknowledgements

N/A

